# Integrative Analysis of *Sorghum Bicolor* Green Prop Roots Under Elevated CO_2_ and Water Deficit Conditions

**DOI:** 10.1101/2021.03.12.435150

**Authors:** Tamires de Souza Rodrigues, Luis Willian Pacheco Arge, Fernanda Alves de Freitas Guedes, João Travassos-Lins, Amanda Pereira de Souza, Jean-Christophe Cocuron, Marcos Silveira Buckeridge, Maria Fátima Grossi-de-Sá, Márcio Alves-Ferreira

## Abstract

Elevated CO_2_ (E[CO_2_]) improves the biomass and yield when combined with water-stress in C4 plants. Although several studies described the molecular response of the C4 plant *Sorghum bicolor* during drought exposure, none reported its combinatorial effect with E[CO_2_] in the roots. We decided to perform a molecular analysis using green prop roots, the portion of the radicular system photosynthetically active and more sensible to drought. Whole-transcriptome analysis identified 394 up- and 1,471 down-regulated genes. Among the E[CO_2_] induced pathways, photosynthesis stood out. Carbon fixation, phenylpropanoid, phenolic compounds, and fatty acid biosynthesis-related pathways were repressed. Protein family analysis showed induction of chlorophyll *a*-*b* binding protein family, and repression of glutathione-related enzymes. Protein-protein interaction networks exhibited well-defined clusters, including genes related to cell organization and biogenesis, oxi-reduction process, and photosynthesis being induced. The findings suggest that the E[CO_2_] mitigates the water deficit by antioxidant and osmoregulation activity, as well as by accumulation of sugar-alcohols in the green prop roots, which may be responsible by the increase in biomass together with the cell proliferation. The higher carbon uptake explains the increase in photosynthetic and primary metabolism activities. Our data revealed that green prop roots present an intriguing metabolism under water deficit and E[CO_2_], showing its crucial role in the drought tolerance acquisition in a predicted future global atmosphere.

## Introduction

Global warming and the increase in atmospheric CO_2_ concentrations ([CO_2_]) are predicted to affect agricultural activities in the near future. CO_2_ emissions contribute to the rising temperatures, which interfere directly with crop productivity due to the intensifying drought events (Musolino et al., 2018). On the other hand, studies describe that elevated [CO_2_] (E[CO_2_]) mitigates the water deficit effects in plants (AbdElgawad et al., 2016; Gustafson et al., 2018; Li et al., 2020).

Although the combined effects of E[CO_2_] and water deficit have been evaluated in soybean and sorghum at the molecular and physiological levels, studies combining these conditions are still scarce in both C3 and C4 plants, (de Souza et al., 2015; Bencke-Malato et al., 2019). Previous works with C4 plants have suggested that E[CO_2_] promotes no physiological alterations in well-watered crop species. However, E[CO_2_] stimulates biomass accumulation and yield under dry growing conditions (van der Kooi et al., 2016). In the C4 plant *Zea mays*, whole-transcriptome analysis identified 1,390 differentially expressed genes in leaf tissues under E[CO_2_] conditions combined with water deficit for 7 and 14 days. Pathways, such as protein phosphorylation and ubiquitination, oxidation-reduction, plant organ development, cellular response to endogenous response, and MAPK signaling, are up-regulated by E[CO_2_] (Ge et al., 2018). De Souza and coworkers (2015) performed a pioneer study in sorghum, using a whole-plant physiologic and metabolomic screening in E[CO_2_] and long-term water deficit, during the grain-filling stage (90 and 120 days after planting, respectively), which is described to be a period more sensitive to drought (de Souza et al., 2015; Prasad et al., 2018). They showed that the sorghum plant adjusted its entire metabolism during drought underCO_2_ exposure. De Souza and co-workers also observed a 94% increase in biomass of the green prop roots (GPR) of drought-treated plants exposed to E[CO_2_] 90 days after planting. The biomass increment in GPRs was 34%, the highest after grains, what is an expressive biomass yield. Moreover, it was reported an increase in sugar-alcohols in the GPR of water-stressed plants (de Souza et al., 2015).

Roots are the first organ to sense the water deficit and respond to it (Hochholdinger, 2016; Zhang et al., 2016; Kreszies et al., 2019). In the radicular system of monocots, prop roots are specific adventitious roots originating from stems above the soil level, which penetrate the soil and anchor the shoot system, preventing them from toppling (Esau, 1977; Glimn-Lacy and Kaufman, 2006). The above-ground portion of prop roots is green and the below-ground portion is white, similar to the rest of root system. Ueno and Fuchikami (2019), by analyzing the sorghum GPR and white portion separately, demonstrated that the morphological structure for both parts is similar, except that GPR present lack of stomata and the presence of granal chloroplasts in the cortex and stele parenchyma (Ueno and Fuchikami, 2019). Experiments conducted with radioactive ^13^C demonstrated that the carbon is transported from leaves to the GPR, explaining the carbon origin and the maintenance of the C3 photosynthesis in an organ without stomata (Ueno and Fuchikami, 2019). Studies with maize and the C4 plant-model *Setaria viridis* reported that prop roots are described to be highly sensitive to drought at physiological and molecular levels, and could be involved in tolerance mechanisms (Zhan et al., 2015; Sebastian et al., 2016). In spite of the importance of GPR for monocots, little is known about its molecular responses to E[CO_2_] and drought stress. Although previous studies described the combinatorial effect of drought and E[CO_2_] in the photosynthesis and growth levels in sorghum plants (Conley et al., 2001; Ottman et al., 2001; Wall et al., 2001; Kakani et al., 2011; Gleadow et al., 2016), as well as the molecular responses under water stress (Fracasso et al., 2016; Varoquaux et al., 2019; Abdel-Ghany et al., 2020), none was reported regarding the molecular effect of E[CO_2_] under water deficit.

The morphological traits, photosynthetic activity and carbon origin in sorghum GPR, as well as its significant biomass increment and sugar-alcohol accumulation observed by De Souza and coworkers (2015) intrigued us to understand the molecular responses of this organ at the transcript level in plants grown in E[CO_2_] under water deficit. Thus, we performed transcriptome analysis of *Sorghum bicolor* GPR at the grain filling stage in E[CO_2_] under water deficit conditions.

## Results

### Whole-transcriptome analysis highlights regulation of the primary and secondary metabolism

GPR samples harvested from control (A[CO_2_] + WD) and treated (E[CO_2_] + WD) groups were used to construct the cDNA libraries. Illumina paired-end sequencing generated between 10.3 and 15.4 million quality reads for *S. bicolor* libraries. Filtered reads percentages were between 87% and 97% (Supplemental Table S2A). The alignment using HiSat showed better results in comparison to TopHat, with the highest average percentage of aligned reads (86.4%; Supplemental Table S2B). Hisat alignment outperformed in differential expression analysis and was used for annotation and enrichment analyses. A total of 1,865 differentially expressed genes (DEGs) were identified, being the majority down-regulated (1,471 genes, 78.87%), and 394 up-regulated (21.13%; Supplemental Table 1).

The *S. bicolor* transcriptome annotation is summarized in Annex I. Based on the GO terms annotated to *S. bicolor* genome from Lawrence-Dill project annotation, we obtained a total of 180,280 GO terms annotated to the 1,865 DEGs (139,528 down- and 40,752 up-regulated). The number of genes for each GO term is summarized in Supplemental Table 1. To better understand the biological information from this dataset, we performed an enrichment analysis.

Enrichment analysis is a statistical approach largely used in transcriptome datasets to identify statistical over/under-represented terms (Khatri et al., 2012). Based on this statistical approach, the Fisher’s exact test from enrichGO function (clusterProfile R package) revealed the following number of GO terms enriched among the down-regulated genes: 276 related to BP (Biological Process), 117 to MF (Molecular Function), and 4 to CC (Cellular Component). On the other hand, the analysis revealed among the up-regulated genes 168 related to BP, 16 to MF, and 15 to CC (Supplemental Table S3A). Aiming to reduce the complexity of GO terms, we focused on the GO terms at level 4, listed in Supplemental Table S3B. We obtained several enriched GO terms, some of which were related to the biological processes of response to stimulus (positive regulation of response to water deprivation: GO: 1902584), among the down-regulated genes. On the other hand, the photosynthesis (chloroplast thylakoid membrane protein complex: GO: 0098807, light-harvesting complex: GO: 0030076, and photosystem: GO: 0009521) biological processes were associated with up-regulated genes.

Analyzing the KEGG pathways, we identified 63 pathways among the down-regulated genes (288) and 31 among the up-regulated ones (78 DEGs) (Supplemental Table S3C). Based on the enrichment analysis, we identified 16 pathways over-represented with DEGs (Supplemental Table S3D; Figure 1). The enriched pathways that were up-regulated included “Photosynthesis-antenna proteins”, “Photosynthesis”, “Glyoxylate and dicarboxylate metabolism”, and “Carbon fixation in photosynthetic organisms”, while “Phenylalanine, tyrosine and tryptophan biosynthesis”, “Amino sugar and nucleotide sugar metabolism”, “Biosynthesis of secondary metabolites”, “ABC transporters”, “Fatty acid biosynthesis”, “Glycerophospholipid metabolism” and “Phenylpropanoid biosynthesis” are the main repressed pathways (Supplemental Table S3D; Figure 1).

**Figure 1.**
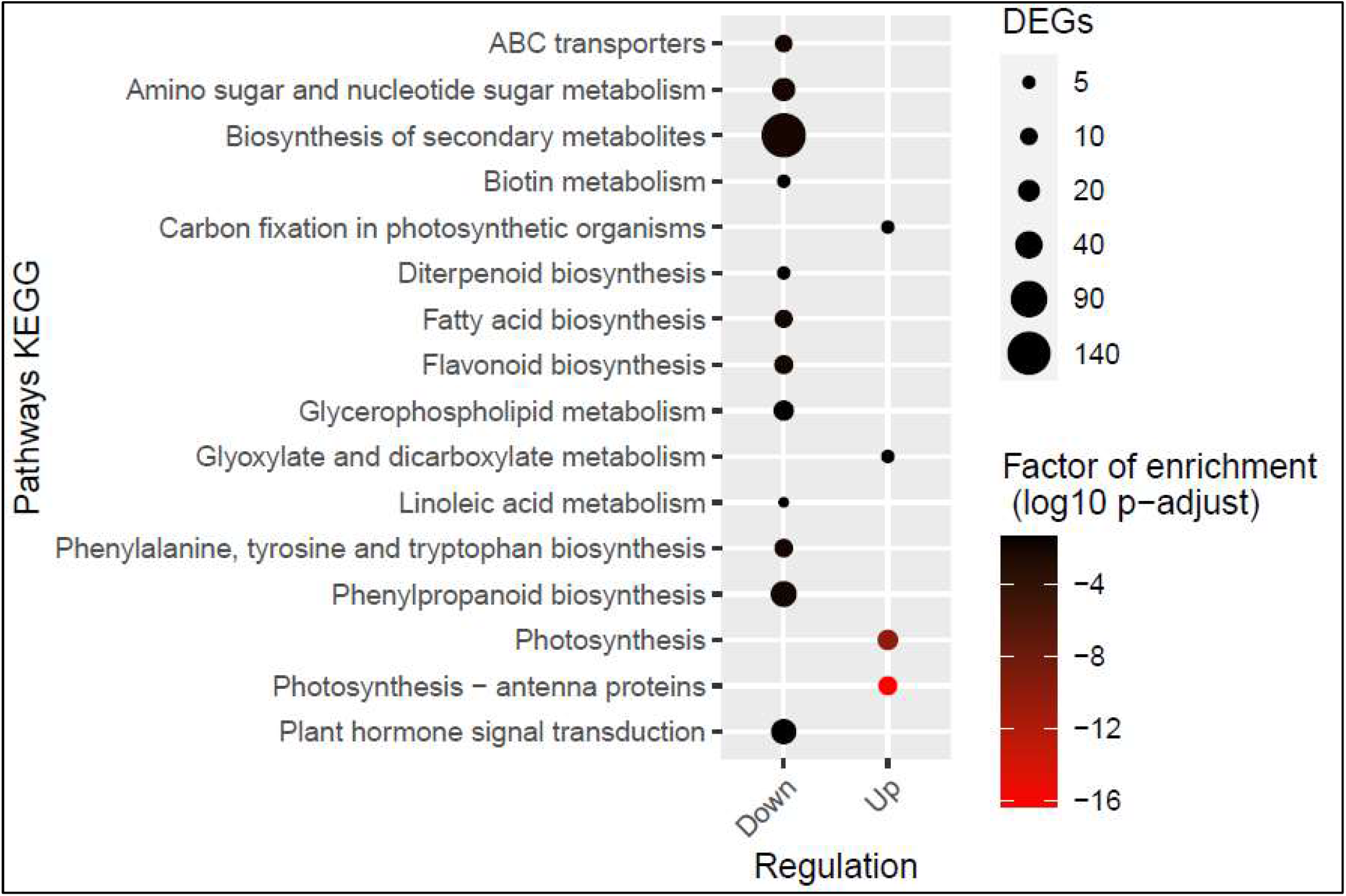
KEGG enrichment analysis. DEGs: differentially expressed genes in response to elevated [CO_2_]. Circle length represents the number of differentially expressed genes (DEGs). The color scale indicates de factor of enrichment (log10 p-adjust).

Based on Protein Family (PFAM) annotation, the enrichment analysis revealed a total of 56 families over-represented among the down-regulated genes, and 17 families among the up-regulated genes (Figure 2). Taking into account the number of DEGs, classes including, “ABC transporter”, “Glutathione S-transferase C-terminal” and “Glutathione N-transferase C-terminal” were significantly down-regulated, while “Cytochrome p450” was up-regulated (Figure 2). According to log10 p-value and the number of DEGs, “Glycosyl hydrolases family 18” and “WRKY DNA-binding domain” were significantly down-regulated. “Chlorophyll A-B binding protein” was up-regulated with the highest log10 p-value (Figure 2).

**Figure 2.**
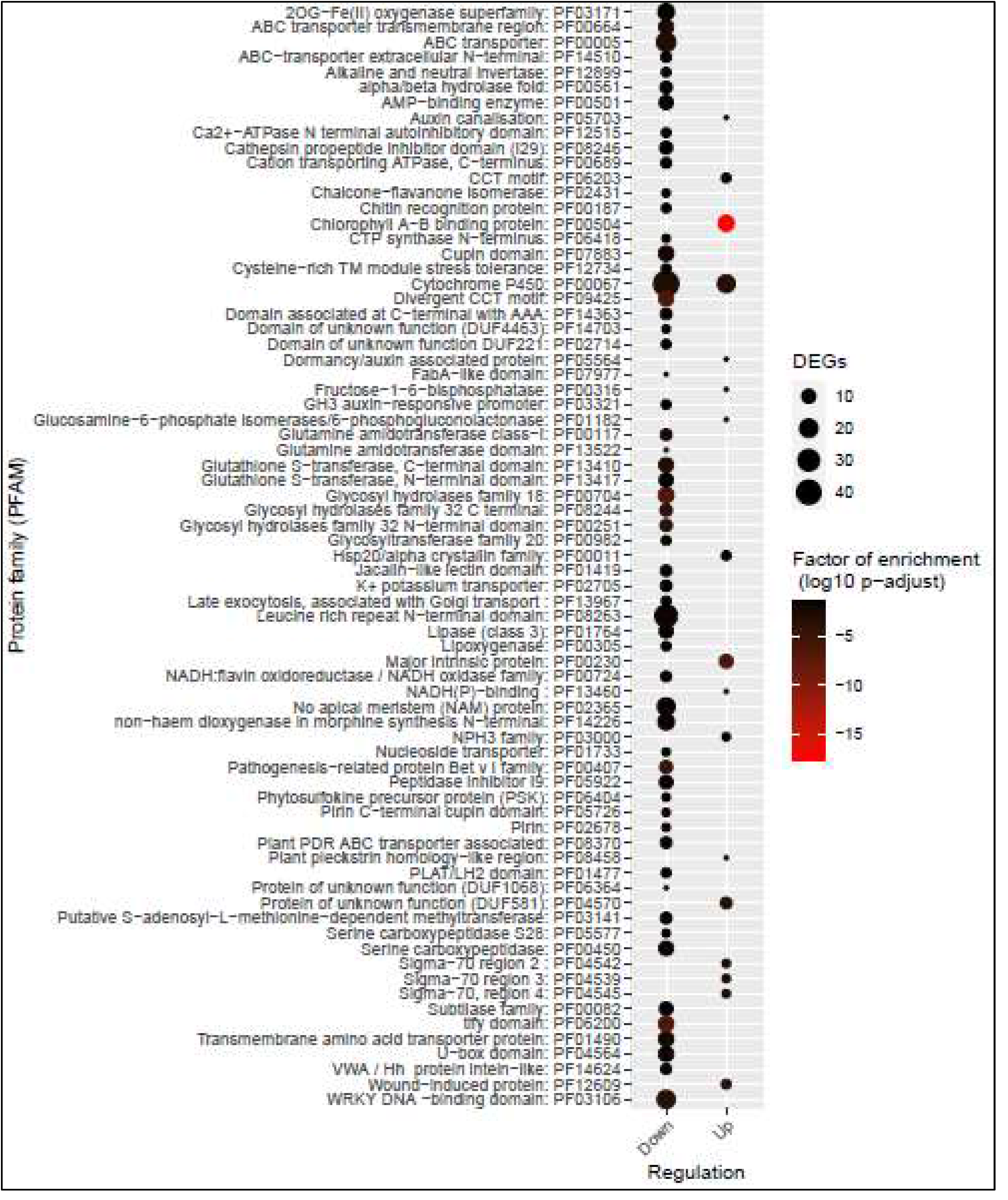
Protein family (PFAM) enrichment analysis.DEGs: differentially expressed genes in response to elevated [CO_2_]. Circle length represents the number of differentially expressed genes (DEGs). The color scale indicates de factor of enrichment (log10 p-adjust).

### Comparative analysis corroborates the modulation of photosynthesis and phenylpropanoid metabolism

Following the GO terms results, the 1,865 sorghum DEGs identified were clustered with the whole-plant (roots, leaves, phloem, and xylem under periodic and chronic drought-heat stress under elevated [CO_2_]) *Populus trichocarpa* study pre-selected. Although poplar is phylogenetically distant from sorghum, it was the only complete RNAseq dataset with a similar experimental design to the one of our study.

The comparison of sorghum genes regulated by elevated [CO_2_] with poplar tissues (Figure 3; Supplemental Table S4) revealed an overlapping group of 148 genes. Among them, sorghum GO terms related to secondary metabolism, as well as induced genes photosynthesis-related were identified (Figure 3; Supplemental Table S4). The genes named light harvesting complex photosystem II subunit 6 (Sobic.006G264201), ferredoxin−NADP(+)−oxidoreductase 1 (Sobic.003G431700), photosystem II light harvesting complex gene B1B2 (Sobic.002G288300), and photosystem II subunit Q−2 (Sobic.002G329600) were down-regulated in both poplar and sorghum roots (Figure 3). Secondary metabolism-related “Pseudouridine synthase/archaeosine transglycosylase-like family protein” (Sobic.006G008700) was down-regulated in GPR sorghum and in xylem poplar tissues (Figure 3; Supplemental Table S4). Regarding phenylpropanoid-related secondary metabolites, “Chalcone and stilbene synthase family protein” (Sobic.005G136300) was repressed in sorghum GPR and poplar leaves, which presented expressive fold-change during chronic stress, and induced in poplar roots (Figure 3; Supplemental Table S4). In addition, it was observed the down-regulation of the “myo-inositol-1-phosphate synthase 2” (Sobic.001G472800) and “Protein kinase superfamily protein” (Sobic.010G200800) genes (Figure 3; Supplemental Table S4). A cluster consisting of the majority of down-regulated genes from sorghum and whole-tissue poplar was represented in the heatmap, in which most of the genes are photosystem-related (Figure 3). Genes named “calcium ATPase 2” (Sobic.007G215200), “PHE ammonia-lyase 1” (Sobic.006G148900), and “amino acid permease 2” (Sobic.009G142800) are also represented in this cluster (Figure 3).

**Figure 3.**
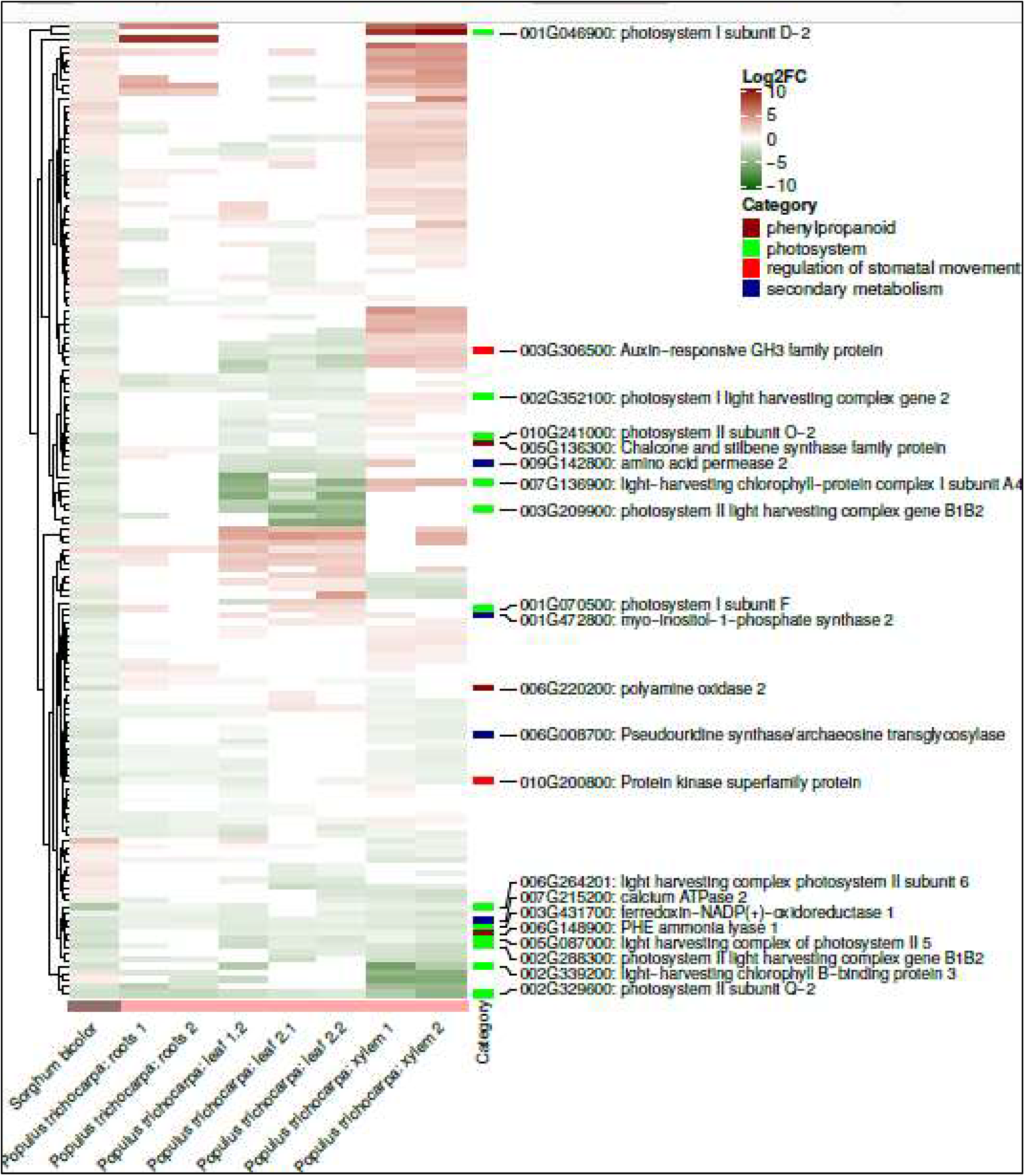
Comparison of *Sorghum bicolor* green prop roots high [CO_2_] responsive genes with *Populus trichocarpa* whole-plant secondary metabolism expression data. The description of the complete data used to construct this heatmap can be found in Table S4.1=periodic stress; 2= chronic stress; leaf 1= young leaf, periodic stress; 2.1=mature leaf, periodic stress; 2.2= mature leaf, chronic stress.

### Integrative system biology highlights the sorghum prop roots mechanisms in response to E[CO_2_]

The protein-protein interaction (PPI) network containing 629 proteins was constructed and resulted in 2,357 interactions (Figure 4A). From the 1,625 DEGs, about 39% were able to form more than one connection in the network. The nodes correspond to the proteins encoded by the DEGs previously identified (Figure 4A). Among the nodes (proteins), 59 were transcription factors (triangles) (WRKY, MYB, NAC, b-ZIP, and bHLH families; Figure 4A). From the clusters formed, which represent a community of nodes highly connected, the two with the highest scores are highlighted in the PPI networks (Figure 4A). Functional enrichment analysis showed that the Cluster I includes 15 nodes and had genes mainly down-regulated and related to the secondary metabolism process, and also related to pigment metabolic process, regulation of biological quality, and oxidation-reduction process (Figure 4A; Supplemental Table S5A). Cluster II contains 70 nodes, which were mainly up-regulated and related to cellular component organization, cellular component biogenesis, oxidation-reduction process, cellular homeostasis, protein folding, and regulation of molecular function (Figure 4A; Supplemental Table S5B). Parameters indicative of network centralities were calculated for each node and the topology scatter plots were performed considering fold-change (Log2FC) and degree (Figure 4B), and fold-change and betweenness centrality (Figure 4C).

**Figure 4.**
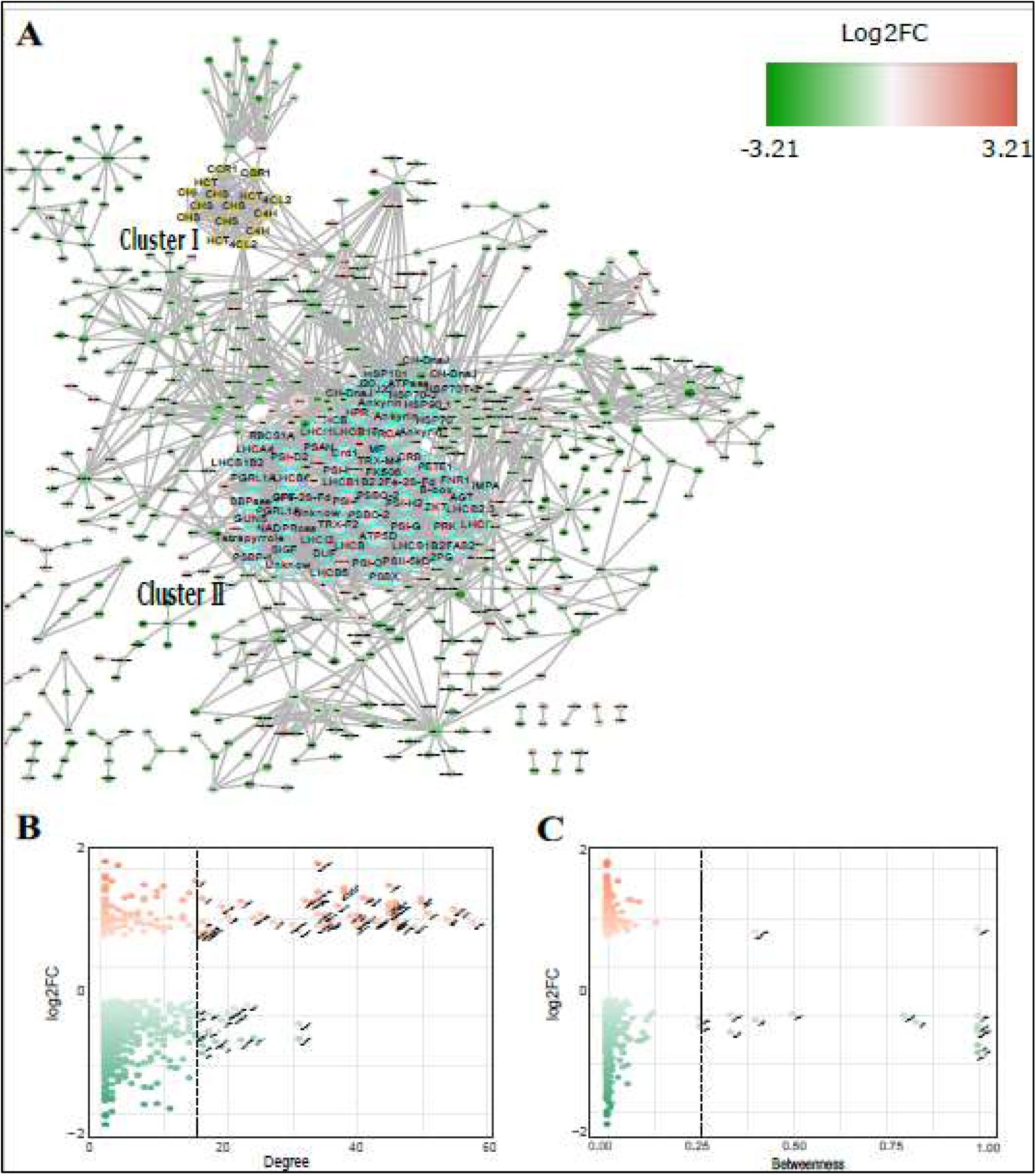
Interactome in response to elevated [CO_2_]. On A, visualization of the protein-protein interaction (PPI) network generated including proteins encoded by differentially expressed genes (DEGs).Clusters were highlighted around the circles in neon blue (Cluster I) and dark blue (Cluster II). On B and C, scatter plots comparing node degree (number of interactions) or betweenness centrality with Log2FC of DEGs. Circles (nodes) represent proteins, while straight lines (edges) represent known or inferred interactions. Circles represent proteins and triangles transcription factors. Red and green represent up- and down-regulated genes, respectively. Gene name abbreviations were included.

The centrality degree measures the number of interactions of the nodes in the network. Up-regulated proteins with high centrality degree were photosynthesis-related, such as light-harvesting complexes (LHC), 2Fe-2S ferredoxin-like superfamily proteins (2Fe-2s) and photosystem I (PSI) subunits, as well as heat-shock proteins. On the other hand, ankyrins, MAP Kinases (MAPK20), cinnamate-4-hydroxylases (C4H) were down-regulated and showed high centrality (Figure 4B). Betweenness centrality measures how frequently the shortest pathway connecting every pair of nodes is crossing a given node [52]. Genes with high betweenness centrality that showed up-regulation were plasma membrane intrinsic protein (PP2) and NAD(P)-linked oxidoreductase superfamily (NADP-LO; Figure 4C); down-regulated genes included some TFs, such as NAC, b-ZIP, ARF and HB (homeobox) families (Figure 4C), as well as proteins related to primary metabolism lipoxygenases (LOX), RNA recognition motif (RRM)-containing protein (RRM), phosphotyrosine protein phosphatases superfamily protein (PFA-PTP) and phytochrome-associated protein 1 (PAP1), and secondary metabolism brassinosteroid-6-oxidase 2 (Figure 4C).

### RT-qPCR shows RNAseq accuration by validation assays

Data of the primers that were selected for RNAseq validation are summarized in Supplemental Table S1. The statistical significance analysis between treated and control-group accessed with REST software confirmed the expression values (Log2FC) obtained by RNAseq. All the tested genes exhibited the same expression pattern, and 19 (95%) were statistically significant (p<0.05). All the thirteen down-regulated genes selected were validated, while six out of seven of the up-regulated genes were statistically validated (p<0.05; Figure S2).

### Metabolome analysis reveals primary metabolism modulation under E[CO_2_]

The metabolites measured by HPLC/MS spectrometry are shown in the Figure S1. The proportion of sugars fructose, trehalose, and deoxyxylulose-5P had a significant increase of about 50% in GPR of plants grown in E[CO_2_] under water-deficit (WD) conditions (p<0.05) in comparison to control-group (A[CO_2_]). All the measured sugar alcohols showed a high level in the treated-group (p<0.01), with inositol increasing 300%, the compound most present in this analysis. The amino acids glutamate, alanine, arginine, aspartate, cysteine, glutamate, histidine, tyrosine, tryptophan, and valine had significant increases (p<0.05). The levels of nucleotides ATP, GDP, CMP, and GMP increased significantly (p<0.05), with values above 50% in treated-groups. The most representative decrease was observed in pentoses-P and the amino acids arginine and ornithine, with values around 50% smaller (Figure S1).

## Discussion

### Assembly of quality S. bicolor green prop roots sequences

In the present study, an average of 13.5 million reads per library was generated, with about 88.5% (12 million, approximately) of reads filtered and 86.4% mapped (about 10.3 million, Table S2A). Previous studies of *S. bicolor* RNAseq using Illumina platform for leaves and grains generated about 47-58 million reads per library and 74.5-86.2% mapped, and about 30,000 genes annotated (Fracasso et al., 2016; Nielsen et al., 2016). Although there was a difference in sequencing depth and the number of annotated genes found in this work, the assigned GO term for the 1,625 annotated genes recovered a great number of GO terms (180,280 GO terms for all DEGs; Table 1). This is due to the recent *S. bicolor* reference genome update (McCormick et al., 2018) available and GO terms annotated by the GOMAP tool, in which the annotation was improved.

**Table 1.** Overview of differentially expressed genes in response to elevated [CO_2_] obtained from functional analysis.

|  |  | **Total** | **Up-regulated** | **Down-regulated** |
| --- | --- | --- | --- | --- |
| **Annotated genes** |  | 1,625 (1,865)<sup>1</sup> | 324 (394)<sup>1</sup> | 1,301(1,471)<sup>1</sup> |
| **GO terms** | Biological process | 124,904 | 28,337 | 96,567 |
|  | Cellular component | 29,676 | 7,327 | 22,349 |
|  | Molecular function | 25,700 | 5,088 | 20,612 |
| **Total** |  | **180,280** | **40,752** | **139,528** |

<sup>1</sup>Numbers between parenthesis represent the total number of genes obtained in the differential expression analysis.

RNAseq is a robust high-throughput sequencing technology with high confidence degree (Marioni et al., 2008). However, the use of RT-qPCR for validating RNAseq is the most acceptable strategy to ensure the accuracy of this technique (Martins et al., 2016). Here, we selected twenty genes to be evaluated by RT-qPCR considering the parameters previously described (see material and methods section). Among these genes, 95% (19) were validated by qPCR, which confirm the accuracy of our RNAseq results (Figure S2). The RNAseq analysis allowed us to identify genes responsive to CO_2_ under water-deficit, constructing a robust integrative analysis between metabolome and transcriptome. The three main pathways regulated include secondary metabolism, primary metabolism, and photosynthesis.

### Elevated [CO_2_] modulates secondary metabolism in GPRs under water-deficit

The enrichment analysis showed a strong modulation of secondary metabolism-related pathways during elevated [CO_2_] (Figure 1) and a large interaction between several proteins related to secondary metabolic, pigment metabolic, regulation of biological quality, and oxidation-reduction process, as observed in the PPI network analysis (Cluster II; Figure 3A; Table S4B). During abiotic stresses, such as drought, salinity, or cold, secondary metabolism has been described to be a key component of the drought tolerance response in plants (Nakabayashi and Saito, 2015). Previous studies have shown that E[CO_2_] modulates plant secondary metabolism, increasing phenolic compounds, monoterpenes and tannins (Peñuelas and Estiarte, 1998; Li et al., 2020). In chickpea, RNA-Seq from 12 tissues in the vegetative and reproductive stages of two cultivars showed an up-regulation of secondary metabolites biosynthesis pathways in E[CO_2_] (Palit et al., 2020b). In soybean roots, secondary metabolism pathways were induced in ambient [CO_2_] in plants water-stressed but no significant alterations regarding gene expression was observed in the combined E[CO_2_] and water-deficit (Bencke-Malato et al., 2019). Beside the phylogenetic distance, sorghum GPR are quite distinct from soybean roots, therefore differences between sorghum and soybean molecular responses are expected since. The secondary metabolism category was considerably down-regulated in sorghum GPR under water-deficit and high [CO_2_] conditions, with 148 DEGs repressed, approximately 8% of the total number of genes regulated (Table S3). It is already well known that E[CO_2_] and drought also induce the production of secondary metabolites (Palit et al., 2020b; Palit et al., 2020a).

Enriched pathways, such as “Phenylpropanoid biosynthesis”, “Phenylalanine, tyrosine, and tryptophan biosynthesis” (phenolic compounds-related) and “Flavonoid biosynthesis” were mostly down-regulated in elevated [CO_2_] during water-deficit (Figure 1). Integrative metabolome analysis showed that the phenylpropanoid biosynthesis pathway had massive down-regulation of several genes, with statistically significant tyrosine accumulation (Figure 5). Palacios (2015) quantified whole-plant phenolic compounds using the same samples used to perform the transcriptome analysis of this study. In GPR under WD, phenylpropanoids were significantly decreased in high [CO_2_], about 6% of the total amount, suggesting that the observed repression of genes related to phenylpropanoid biosynthesis is directly associated with the phenylpropanoids depletion (Jara, 2017). For instance, cinnamate-4-hydroxylase (ID: Sobic.003G116800, Annex I), the first Cytochrome P450-dependent monooxygenase of the phenylpropanoid pathway (Bell-Lelong et al., 1997), is down-regulated and it can be seen in the PPI network analysis (Figure 3B).

**Figure 5.**
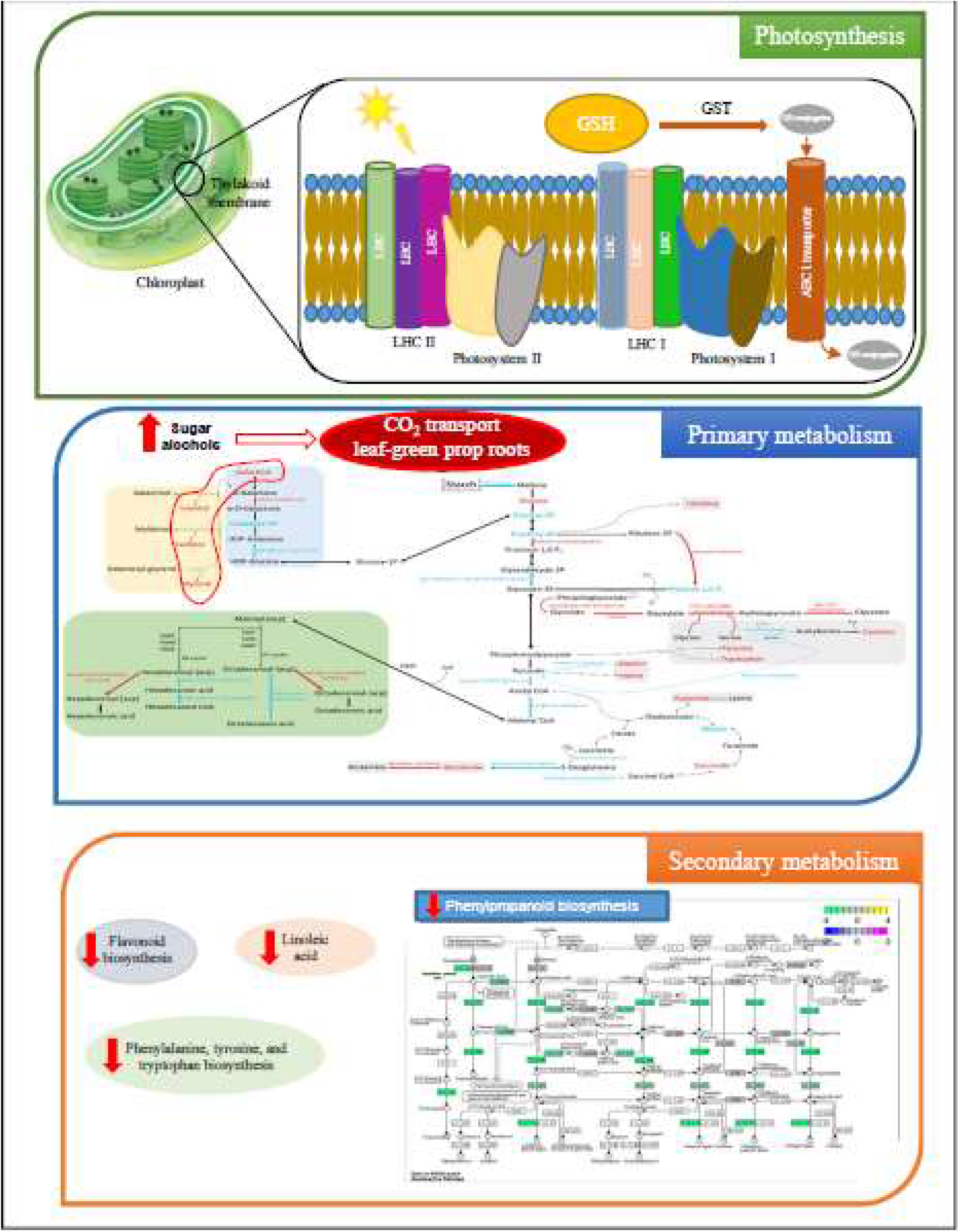
Overview of the main pathways regulated in *Sorghum bicolor* green prop roots transcriptome. Genes related to photosynthesis, primary and secondary metabolism were modulated.

E[CO_2_] has been associated with repression with secondary metabolites production. In strawberry fruits, elevated [CO_2_] inhibited the phenylpropanoid and flavonoid pathways, identified through the evaluation of key-genes by RT-qPCR (Li et al., 2019). Regarding metabolites, the comparison between two variants from *Oryza sativa* reduced the total content of phenylpropanoids and flavonoids in plants exposed to 550 ppm of CO_2_ (Goufo and Trindade, 2014), showing that a slight alteration in CO_2_ concentration was able to modify the amount of these phenolic compounds. In contrast to the study mentioned above, durum wheat leaves exposed to E[CO_2_] in well-watered conditions showed in transcriptome analysis an up-regulation of 66.6% of the phenylpropanoid-related pathways (Vicente et al., 2019).

Phenylalanine, tyrosine, and tryptophan-enriched biosynthesis pathways are directly related to the phenylpropanoid biosynthesis route, because of the repression cascade of all genes of the phenolic compounds-related identified in our analysis (Figure 1). It can be explained by the origin of the phenolic compounds biosynthesis pathways. These three amino acids are originated from the shikimic acid pathway in the primary metabolism of vascular plants (Buchanan et al., 2015). The key enzyme of the balance between phenylalanine, tyrosine, and tryptophan production and the first secondary metabolism compounds is phenylalanine ammonia-lyase (PAL), which converts phenylalanine in trans-cinnamic acid by a non-oxidative reaction (Camm and Towers, 1973). Phenylalanine is the primary substrate for the phenylpropanoid pathway that gives rise to flavonoids and anthocyanins, important antioxidant molecules that act to protect plants of the oxidative stress promoted during water-deficit (Vojta et al., 2016). Our results showed a down-regulation in E[CO_2_] of *PAL1* (Sobic.006G148900; Annex I), significantly increased levels of the amino acids tyrosine and tryptophan and no significant alteration in phenylalanine metabolite levels (Figure S1). Furthermore, *PAL1* poplar homologs were also repressed in leaves under drought and E[CO_2_] (Figure 3; Table S4), exhibiting similar behavior to sorghum GPR.

Our results of the pathway enrichment analysis showed the repression of the linoleic acid metabolism pathway in response to elevated [CO_2_] under water deficit (Figure 1). Roots submitted to short-term water-deficit and elevated [CO_2_] in soybean also presented a down-regulated *LOX1* (Lipoxygenase) gene (Bencke-Malato et al., 2019). In particular, thr enrichment analysis identified a down-regulation of linoleate 9s-lipoxygenase (*LOX1_5*; ID: Sobic.001G483400; Annex I) and interactome analysis reported *LOX3* as repressed in PPI network (Sobic.005G054900; Annex I; Figure 3A and C). LOXs are enzymes that catalyze the oxygenation of polyunsaturated fatty acids, such as linoleic and linolenic acids (Gong et al., 2015), and are converted into a variety of bioactive mediators involved in biotic and abiotic stresses, even though few studies have been carried out in plants to establish the role of LOX during abiotic stress (Viswanath et al., 2020). *CaLOX1* overexpression in pepper promoted high expression levels of oxidative response-related genes, as well as reduction of H_2_O_2_ accumulation and lipid peroxidation during drought stress (Lim et al., 2015). In finger millet seedlings, water-deficit increased total LOX activity and lipid peroxidation (Kotapati et al., 2014). Therefore, we showed the repression of a high number of genes related to secondary metabolism pathways, specifically down-regulation of phenylpropanoid and whole-phenolic metabolism pathways, as well as the depletion of their metabolites in response to E[CO_2_].

### Elevated [CO_2_] modulates primary metabolism and cell biogenesis under water-deficit

Genes related to primary metabolism showed an important role in sorghum response to E[CO_2_]. Two enzymes related to Calvin-cycle (phosphoribulokinase and fructose-1,6-bisphosphatase) were up-regulated in the carbon fixation enriched pathway in our experiment. In addition, PPI network analysis identified 27 proteins (39% of total genes identified in the Cluster I) related to cell component biogenesis and organization with high degree centrality and mainly up-regulated (Figure 3A; Table S4A). In *Ipomoea trifida* roots, drought promoted down-regulation of most of the genes related to cell component, biogenesis and organization (Zhu et al., 2017). A study performed with distinct grasses revealed reduction on number of crown (or prop) roots and suppression of their development during water-stress, as well as the GO terms related to cell component and cell wall biogenesis were down-regulated in *Setaria viridis* crown roots (Sebastian et al., 2016). In general, root growth of crop species is stimulated by E[CO_2_] and this is well elucidated (ROGERS et al., 1992; Wechsung et al., 1999; Pritchard et al., 2006; Madhu and Hatfield, 2013; Bahrami et al., 2017; Uddin et al., 2018), even though the combined effect of water-deficit and E[CO_2_] is not clearly involved in the root development of the C4 plants. However, based on our analysis we suggest the E[CO_2_] may contribute to an increase in cell division and growth, and consequently biomass accumulation in sorghum GPR. Apparently, the E[CO_2_] is also intensifying the activation of genes related to glucose and fructose production in drought conditions. The metabolome analysis showed that glucose content increased significantly in E[CO_2_], as well as all sugar-alcohols quantified (Figure S1). In wheat grains, metabolome response to [CO_2_] showed an increase in glucose, fructose, sugar alcohols such as myo-inositol and glycerol-2P (Högy et al., 2010), while in stolon of creeping bentgrass [CO_2_] caused an increase in the relative content of eight sugars (including fructose), glycerol, and myo-inositol sugar alcohols (Xu et al., 2018). Sugar-alcohols (or polyols) are highly soluble and chemically inert sugars originated from the reduction of sugar, and they present different functions as osmoprotective molecules under abiotic stresses (Singh and Laxmi, 2015). Together with sucrose, these soluble sugars are responsible for transport and storage of carbon via phloem for energy and carbon supply (Jain et al., 2010). Furthermore, up to 30% of the carbon fixed by plants are in the form of polyols (Bieleski, 1982). Thus, the regulation of genes related to sugar-alcohols metabolism, as well as its significant metabolite accumulation suggest that the E[CO_2_] intensifies the osmoprotective responses, and that it is a relevant carbon uptake route participating in the carbon transport from leaves to GPRs. Fatty acid (FA) and glycerophospholipids biosynthesis pathways were regulated in E[CO_2_]. *Sorghum* roots transcriptome analysis previously performed showed FA metabolism and biosynthesis pathways enriched under osmotic stress promoted by salt (Yang et al., 2018), but no studies evaluating the FA in combined drought and E[CO_2_] conditions were performed so far. Interestingly, all saturated FA-related genes are down-regulated. On the other hand, a chloroplastic key enzyme involved in the biosynthesis of unsaturated FAs in plants (Behrouzian and Buist, 2002) named stearoyl-[acyl-carrier-protein] 9-desaturase 1 is up-regulated (gene ID Sobic.003G404000; Annex I). Although the total fatty acid content presented no difference between ambient and E[CO_2_] treatments under water-deficit in GPR (de Souza et al., 2015) at metabolomic level, the transcriptome indicated repression of saturated FA pathway (Figure S3). On the other hand, unsaturated FA pathway is up-regulated in GPR under drought and E[CO_2_]. (Figure S3). Unsaturated FAs constitute complex lipids that are essential components of cellular membranes, being a source of energy stored as triacylglycerols (Hernández et al., 2019). Moreover, the FA degradation pathway was also repressed (data not shown), suggesting that the metabolism is being directed to unsaturated FAs production.

In addition, we identified the down-regulation of glycerophospholipid metabolism in GPR (Figure 1). No study so far has reported the molecular role of the glycerophospholipids in roots transcriptional response, but Li and coworkers described the same pattern in cucumber seedlings leaves in response to elevated [CO_2_] under moderate drought stress (Li et al., 2018).

Therefore, we suggest that the up-regulation of genes related to carbon fixation and cell biogenesis and organization, glucose accumulation, as well as the increase in unsaturated fatty-acid transcripts and sugar-alcohol metabolites are, together, involved to the increase of biomass in prop roots of plants grown at E[CO_2_] under drought stress.

### Carbon uptake in E[CO_2_] stimulates GPR photosynthesis under water-deficit conditions

Enrichment and the protein family analyses showed the induction of photosynthesis-related pathways and genes involved in the photosynthetic electron transport chain (ETC) (Figure 1 and 2). In well-watered conditions, E[CO_2_] contributes to stomatal closure during long-period exposure because of the saturation of the cell CO_2_ scavenge system (Gamage et al., 2018). Besides, glutathione is considered the most important antioxidant defense during drought stress and acts as the substrate for glutathione S-transferase (GST) (Aquilano et al., 2014). Four genes encoding GSTs were highly up-regulated at 1h of water-deficit treatment in sorghum (Abdel-Ghany et al., 2020), but we did not find any report on glutathione metabolism-related in response to E[CO_2_]. Here, the GST domain protein families are down-regulated in combined water deficit and E [CO_2_] (Figure 2) condition, as well as the ABC transporters pathway (Figure 1) and ABC transporter protein families-related (Figure 2). ABCC1 transporter was previously identified in sorghum and are pumps for glutathione S-conjugates in the thylakoid membrane from chloroplast (OJ and CG, 2018).

Among the ETC-related genes, the Psb S subunit of the photosystem II (PSII) and light-harvesting chlorophyll protein complex (LHC) in the antenna complex were significantly up-regulated. In addition, the family of the chlorophyll *a*-*b* binding proteins was predicted to be down-regulated (Figure 2). These proteins belong to the LHC and are responsible for the light scavenging in the PSII (Liu et al., 2013), which explains it being also repressed in E[CO_2_]. Hence, our findings indicate the [CO_2_] acts mitigating the effects of oxidative damage in the photosynthetic apparatus caused by water deficit through the reduced expression of ROS scavenging via GSH metabolism and chlorophyll *a*-*b* binding protein, as well as progressive activation of ETC-related genes.

## Conclusions

The large amount of pathways regulated in sorghum GPR indicates its wide metabolic activity under E[CO_2_] and water stress. The summary of the main GPR pathways regulated by E[CO_2_] under water deficit is shown in Figure 6. Indeed, the molecular response to E[CO_2_] indicated its alleviate upon WD in sorghum GPR, as described in other species (Xu et al., 2015; AbdElgawad et al., 2018; Li et al., 2018; Li et al., 2020; Wu et al., 2020). The repression of secondary metabolism and antioxidant activity by GSH biosynthesis pathway activation during the photochemical photosynthesis phase showed the GPR decreased the energy investment in plant defense, which was probably driven towards photosynthesis and biomass accumulation (Figure 6).

The particular transcriptional and metabolomic networks of this organ are clearly associated with the significant biomass yield previously reported (de Souza et al., 2015). Our initial hypothesis was that this gain of dry-weight could be associated with complex-chain carbohydrates, such as lignin and/or cellulose. However, the results showed that the carbon fixation is directed towards glucose synthesis and the production of high amounts of sugar-alcohols (polyols). Although this discovery seems incoherent, it makes sense if we consider that the polyols are soluble sugars, being the main vehicle for carbon transport in GPR under high [CO_2_]. Due to this, we hypothesize that the excess of CO_2_ is transported as polyols, displaying a central role in the carbon translocation from the leaves to GPR (Figure 6), as well as in the induction of gene expression related to cell biogenesis, which can explain the biomass accumulation observed in this organ (de Souza et al., 2015). Ueno (2019) described this CO_2_ transport, even though they did not identify the mechanism behind this flux. Besides, this type of transport via polyols was previously described in plants (Uddin et al, 2018). Then, the soluble sugar accumulation could mean this organ works as an immediate energy source for the plant in high [CO_2_], exhibiting a relevant drought tolerance mechanism. Furthermore, the photosynthetic activity stimulated by the upper carbon availability contributed for the glucose production, also being an important energy source.

## Material and Methods

### Plant material and experimental conditions

*Sorghum bicolor* BRS330 was cultivated in soil, as described by De Souza et al. (2015). Plants received 1.5 L of water per day during the vegetative stage (0 - 60 days after planting, DAP). When the reproductive stage started (after 60 DAP), water deficit treatment was performed by reducing the irrigation to 0.45 L of water per day (70% of field capacity reduction). Control-plants were grown in ambient [CO_2_] (A[CO_2_], 400 µmol mol^-1^), and treated-plants in elevated [CO_2_] (E[CO_2_], 800 µmol mol^-1^), both submitted to chronic water-deficit (WD) for 30 days. Green prop roots (GPR) were harvested at 90 DAP, which corresponds to the grain filling stage when the grains are immature. Harvested samples were stored at room temperature (25°C) for metabolome and at −80°C for transcriptome analyses (de Souza et al., 2015).

### Metabolites quantification

Metabolites were extracted after lyophilization of the samples maintained at ambient temperature. The quantification of intracellular metabolites was performed with LC-MS/MS (UHPLC device from Agilent Technologies/QTRAP 5500; AB Sciex). Methodology details are available in De Souza et al. (2015). Statistical analysis between control and treated groups was performed using the Kruskal-Wallis test, followed by Dunn’s test, for multiple comparisons, in the GraphPad Prism 6 software. Results were expressed as mean ± SD (standard deviation). Values with p ≤ 0.05 were considered significant.

### RNA extraction, cDNA library construction and Illumina sequencing

Total RNA extraction was performed with 100 mg of grounded GPR tissue using the ReliaPrep RNA Tissue to fibrous tissues (Promega). The RNA quality and purity were accessed with NanoDrop 2000 Spectrophotometer (Thermo Scientific). Samples with 260/280 and 260/230 ratios between 1.7 and 2.1 were considered acceptable for further analysis. RNA quantification on Qubit fluorimeter (Thermo Fisher Scientific) varied between 53.4 and 133.0 ng µL^-1^. RNA integrity was verified in Bioanalyzer 2100 (Agilent Technologies) using Agilent RNA 6000 Pico Kit. Samples with high quality (RIN > 7) values were obtained. Eight biological replicates (four for each control and treated-groups) containing one plant per sample were selected for RNA sequencing. 1 µg of total RNA was used to synthesize paired-end (2×100 bp) first-strand libraries, which was constructed according to the manufacturer’s protocol of the Truseq Stranded mRNA Sample Prep LS Kit (Illumina). The 16 first-strand paired-end (2×100 bp) libraries (8 biological x 2 technical replicates) were sequenced using an Illumina HiSeq 2500 platform. The mRNA library construction and sequencing were carried out in the Brazilian Bioethanol Science and Technology Laboratory (CTBE, Campinas-SP, Brazil). Sequences is being deposited in NCBI (accession number xxxxx).

### Data preprocessing and alignment

The preprocessing step was carried out with Trimmomatic v0.36 (Bolger et al., 2014) to remove adapters and filter raw reads by quality. Since the sequencing depth was not so high, a smooth read preprocessing approach using the Trimmomatic-suggested parameter values was preferred. The filtered quality reads were aligned to the most recent version (v3) of the *Sorghum bicolor* genome downloaded from the Phytozome database (http://www.phytozome.net) using the Hisat2 software (Pertea et al., 2016) with its default parameter values.

### Differential expression analysis

The aligned read counts, the fragments per kilobase of transcript per million mapped reads (FPKM) normalization values, and the differential expression analysis were carried out using the Cuffdiff package from Cufflinks v2.2.1 (Trapnell et al., 2012) and its default parameter values. Genes with FDR value < 0.05 were considered differentially expressed genes (DEGs).

### 2.6 Functional annotation and enrichment analysis

Annotation information of the genes was retrieved from different databases: gene function, protein family (Pfam) (Finn et al., 2014) and KEGG pathways (Kanehisa, 2000) from the Phytozome database; transcription factor (TF) from PlantTFDB database (Zhang et al., 2011), high coverage annotation Gene Ontology (GO) (Ashburner et al., 2000) terms from the project Gene Ontology Meta Annotator for Plants - GOMAP (Lawrence-Dill, 2019). To use the Pathview R package (Luo and Brouwer, 2013), which is used to plot KEGG pathways and is based on the NCBI annotation, we performed a conversion of IDs (Phytozome to Entrez) by identifying the best hit between them via local protein BLAST analysis (Camacho et al., 2009), with the following parameters: identity ≥ 70% and e-value < 1e-10.

To identify relevant annotation factors in the differentially expressed genes (DEGs) dataset, we performed an enrichment analysis applied to PFAM, TF, GO terms, and KEGG pathways. To identify overrepresented PFAM and TF annotation factors and KEGG pathways, we applied a Fisher’s exact test (Huang et al., 2009), followed by the correction of all p-values with the False Discovery Rate (FDR) method (Noble, 2009). The enrichment analysis of GO terms was performed with enrichGO function from the clusterProfiler R package (Yu et al., 2012) and then submitted to a filter at the level 4 with gofilter function from GOSemSim R package (Yu et al., 2010). We adjusted all p-values with the Bonferroni-Hochberg method. Each enrichment analysis was performed with the DEG dataset separated in up and down-regulated genes, compared to all annotated genes of the genome, and annotation factors with adjusted p-value < 0.05 were considered as enriched. For graphical representation purposes, the adjusted p-values were transformed using log10 scale to represent the level of enrichment. The ggplot2 R package (Wickham, 2016) was used to construct bar and bubble charts.

### Comparative analysis between S. bicolor GPR and Populus trichocarpa

A survey for related studies in the literature (E[CO_2_] and water stress) was performed in order to compare the sorghum GPR expression pattern to *Populus trichocarpa* (Georgii et al., 2019), *Glycine max* (Bencke-Malato et al., 2019), *Oryza sativa* (Shen et al., 2011), and *Zea mays* (Opitz et al., 2014). After the selection of the studies, we performed the best hits identification between *S. bicolor* and each other species via Blastp analysis based on the following thresholds: percentage of identity ≥ 70% and e-value < 1e-10. Based on the best-hit identification we built a heatmap using the R package Complex Heatmap Ver. 2.5.1 (Gu et al., 2016), to compare the distinct profiles. Only the *P. trichocarpa* dataset shared a considerable number of genes with our data. Therefore, we decided to represent these two species.

### Systems biology approach

To study the DEGs in a holistic way, we used an approach from the systems biology field. We used a protein-protein interaction (PPI) approach to aid in the study of the complexity of biological systems that form the whole of living organisms. In those, the system is more than the sum of their parts. All interactions between proteins from *S. bicolor* related to the DEGs dataset of this study were downloaded from STRING Database v.12. To get the PPI of *S. bicolor*, we converted the gene identifiers from v.3 to v.2, based on the Phytozome annotation file. The criteria selected to build the network considered the E[CO_2_] experiments, databases, and co-expression, with threshold > 0.2.

The network was built in Cytoscape software v.3.7.2 (Shannon, 2003), the topology analyses of degree and betweenness centrality were performed with the Network Analyzer (Assenov et al., 2008), and the module detection was performed with MCODE v.1.6 (Bader and Hogue, 2003). The topology centralities results were exported and then used to build scatter plots in R with the ggplot2 package. Modules with size greater than 10 nodes were used in an enrichment analysis of GO terms (FDR<0.05), performed with a custom R script using FDR < 0.05 to consider overrepresented GO terms.

### RNAseq validation by RT-qPCR

Twenty genes were selected for RNAseq validation, considering Log_2_FC ≥ ±1.5 and FPKM > 100. Two reference genes were selected based on a previous study made with *S. bicolor* under water-deficit conditions (Reddy et al., 2016). For primer design, Primer3Plus (http://bioinformatics.nl/cgi-bin/primer3plus/) was used considering the following parameters: melting temperature around 60°C, GC content between 40-60%, primer size between 19-20 bp and amplicon length lower than 250 bp. The amplicon length varied from 108 to 245 bp (Table S1). Total RNA (1µg) extracted was used for cDNA synthesis using SuperScript™ III Reverse Transcriptase (Invitrogen) reaction, according to the manufacturer’s instructions. The reaction (20 µL) was incubated at 50°C for 1h followed by enzyme inactivation at 70°C for 15 min. PCRs with specific primers were performed and bands observed in 1% agarose gel electrophoresis to identify gDNA contamination. cDNA samples were diluted 50 times prior to use in RT-qPCR assays. The melting curve analysis of the amplification products confirmed that the primers amplified only a single product with the expected size (data not shown).

RT-qPCR was performed in a 96-well plate on the 7500 Fast Real-Time PCR System (Applied Biosystems). The cDNA amplification reactions contained: 2 μL of the SYBR Green detector (1x) (Molecular Probes); 0.4 μL of ROX dye; 10 μL of 1:50 diluted cDNA; 0.8 μL of the respective primer pair (10 μM each); 0.05 μL of each dNTP (10 mM); 2 μL of PCR buffer (10x); 1.2 μL of 50 mM magnesium chloride; 0.25 units of Platinum® Taq DNA Polymerase (Invitrogen™); and 3.5 μL of water (final volume: 20 μL). In the control reaction, the cDNA was replaced by ultrapure and sterile water. The reaction parameters included initial denaturation at 94°C for 5 min, followed by 40 cycles at 94°C for 15 s, annealing of primers at 60°C for 10 s, extension by the enzyme at 72°C for 15 s and fluorescence reading at 60°C for 35 s. The specificity of the amplification was analyzed through the dissociation curve profiles. Assays were performed using three technical replicates. Quantification cycle (*Cq*) and primer efficiency were calculated by Miner online tool (Zhao and Fernald, 2005). qBase v1.3.5 software was used to calculate the non-normalized expression values, using the *Cq* values (Hellemans et al., 2007). The relative expression level (Log_2_FC) and statistical analyses between control and treated groups were performed on the Relative Expression Software Tool (REST) (Pfaffl et al., 2002).

## Acknowledgements

We would like to thank Dr. Douglas Paixão and Jean-Christophe Cocuron for sample preparations for RNA sequencing and LC/MS analyses, respectively. RNAseq was performed in the Brazilian Bioethanol Science and Technology Laboratory (CTBE), Campinas, SP, Brazil, and LC/MS in the Center for Applied Plant Sciences, Ohio State University, Columbus, USA.

## Notes

This research was supported by the CNPq (National Council for Scientific and Technological Development), CAPES (Coordination for the Improvement of Higher Education Personnel/ Ministry of Education) and FAPERJ (Carlos Chagas Filho Foundation of Support for Research in the State of Rio de Janeiro). This work is part of the doctoral thesis of Tamires S. Rodrigues in Plant Biotechnology (Programa de Pós- Graduação em Biotecnologia e Bioprocessos, Universidade Federal do Rio de Janeiro, Brazil), supported by a scholarship from CNPq (grant number 162676/2015-8). The author’s research was supported by Conselho Nacional de desenvolvimento Científico e Tecnológico (CNPq) (grant number 308832/2017-5), Instituto Nacional de Ciência e Tecnologia (INCT Biotec Seca-Pragas (grant number 465480/2014-4) and INCT-Bioethanol (FAPESP2008/57908-6 and CNPq 574002/2008-1)), and Fundação de Amparo à Pesquisa do Rio de Janeiro (FAPERJ CNE; grant number E-26/202.631/2019).

